# PrP^C^-induced signaling in human neurons activates phospholipase Cɣ1 and an Arc/Arg3.1 response

**DOI:** 10.1101/2025.04.01.645054

**Authors:** Daniel Ojeda-Juarez, Gail Funk, Emily Richards, Alexander J. Rajic, Daniel B. McClatchy, Katrin Soldau, Xu Chen, John R. Yates, Steven L. Gonias, Christina J. Sigurdson

## Abstract

Synaptic dysfunction and loss correlate with cognitive decline in neurodegenerative diseases, including Alzheimer’s disease (AD) and prion disease. Neuronal hyperexcitability occurs in the early stages of AD and experimental prion disease, prior to the onset of dementia, yet the underlying drivers are unclear. Here we identify an increase in the immediate early gene, Arc/Arg3.1, in the human prion disease-affected frontal cortex, suggestive of neuronal hyperactivity. To investigate early signaling events initiated by prion aggregates (PrP^Sc^) in human neurons, we stimulated PrP^C^ in human iPSC-derived excitatory neurons (iNs) with a known PrP^Sc^-mimetic antibody (POM1), which recapitulated the Arc/Arg3.1 response within two hours. Proteomics, RNAseq, and a phosphokinase array in iNs revealed alterations in the EGF receptor and increased phosphorylated phospholipase C (PLC)-γ1 (Y783), which was also observed in the cerebral cortex of prion-infected mice. Thus, PrP^C^ ligands can induce a PLC-γ1 intracellular signaling cascade together with an Arc response, suggestive of a neuronal activity response.

## Introduction

Synapse loss and neuronal hyperactivity occur early in Alzheimer’s disease before pronounced neuronal loss and hypoactivity are evident (1, 2). Similarly, synapse loss is an early neuropathological feature of prion-induced neurodegenerative disease (3–8), and the onset of clinical signs correlates with reduced synaptic proteins (5, 9). We previously found that prion-infected mice show an early and persistent increase in Arc/Arg3.1 (activity-regulated cytoskeleton-associated protein), an immediate early gene responsive to enhanced neuronal activity (10). The pronounced long, concave post-synapses observed in prion-infected brains are also suggestive of heightened synaptic activity (9, 11–13). Yet the mechanisms underlying dysregulated neuronal activity and Arc/Arg3.1 (hereafter referred to as Arc) induction in prion disease, and how human neurons respond to PrP^C^-induced signaling are unknown.

*Arc* is rapidly induced by either a rise in intracellular Ca^2+^ or an increase in cyclic adenosine monophosphate (cAMP) (14, 15), thus serving as an early indicator of increased glutamatergic activity. Arc functions as a master regulator of synaptic plasticity, promoting long-term depression (LTD) by regulating synaptic clustering of α-amino-3-hydroxy-5-methyl-4-isoxazolepropionic acid receptors (AMPARs) (16). Arc competes with PSD95 for binding to transmembrane AMPA receptor regulatory proteins (TARPs), with TARP-bound Arc resulting in AMPAR dispersal from the postsynaptic density (PSD) condensate, facilitating endocytosis in weaker synapses (17–19) [“inverse” synaptic tagging (20)].

Prion disease is characterized by the conversion of the cellular prion protein (PrP^C^) to a pathological β-sheet-rich misfolded form, PrP^Sc^ (21, 22). PrP^C^ binds prion oligomers (PrP^Sc^), as well as amyloid-β (Aβ), tau, and α-synuclein oligomers, which results in the activation of macromolecular complexes and signaling at the post-synapse (23–27). However, as a glycosylphosphatidylinositol (GPI)-anchored protein, PrP^C^ lacks an intracellular domain and thus requires a transmembrane partner for intracellular signaling. PrP^C^ has been reported to interact with a variety of cell surface receptors, including epidermal growth factor receptor (EGFR), G protein-coupled receptors (GPCRs), low-density lipoprotein receptor-related protein-1 (LRP1), neural cell adhesion molecule 1 (NCAM1) and N-methyl-D-aspartate receptors (NMDARs) (24, 28–31). Although multiple receptors may interact with PrP^C^, relatively little is known about how these interactions regulate cell signaling in prion disease, particularly in the context of a human neuron.

Studies using human induced pluripotent stem cells (iPSC)-derived cells or organoids to model human diseases have focused on genetic prion disease and reported aberrant synaptic structure, excitatory-inhibitory imbalance, altered bioenergetics (32–37), and the ability of human cells to replicate prion oligomers *in vitro* (38–40), however, these chronic disease models have not elucidated the acute cell-signaling events that may occur within hours of prion exposure. Herein, we discovered increased Arc in patients with familial or sporadic prion disease. To begin to understand how early PrP^C^-triggered cell signaling events impact Arc, we stimulated human iPSC-derived excitatory neurons (iNs) with a PrP^Sc^-mimetic antibody (POM1) (41, 42), and used proteomics, RNA-seq, and phosphokinase arrays to identify downstream cell-signaling events that occur by two hours. Our results recapitulate the increase in Arc observed in prion disease, and reveal an increase in phosphorylated phospholipase C (PLC)-γ1 and ERK1/2, and a decrease in AKT and S6 kinase phosphorylation. PrP^Sc^ also induced higher phospho-PLC-γ1 in the brain of prion-infected mice. This human-relevant neuronal model reveals the PrP^C^ - PLC-γ1 axis as an acutely activated signaling network in prion disease, as well as a potential therapeutic target.

## Results

### Arc is increased in human sporadic and genetic prion disease

We first measured Arc as a proxy for neuronal activity in the frontal cortex of patients with sporadic CJD (sCJD) (n = 6) and Gerstmann-Sträussler-Scheinker (GSS) (n = 4) patients [GSS: familial prion disease; P102L (n = 1) and F198S (n = 3) in the *PRNP* gene] by western blot. Frontal cortex is commonly altered in prion-affected brain by magnetic resonance imaging (MRI) (43, 44). sCJD and GSS brain samples showed an approximately 4 – fold and 1.25 – fold increase in Arc, respectively, compared to age-matched controls (Fig. 1A – B; Table 1), suggestive of heightened neuronal activity in human prion disease.

**Figure 1.**
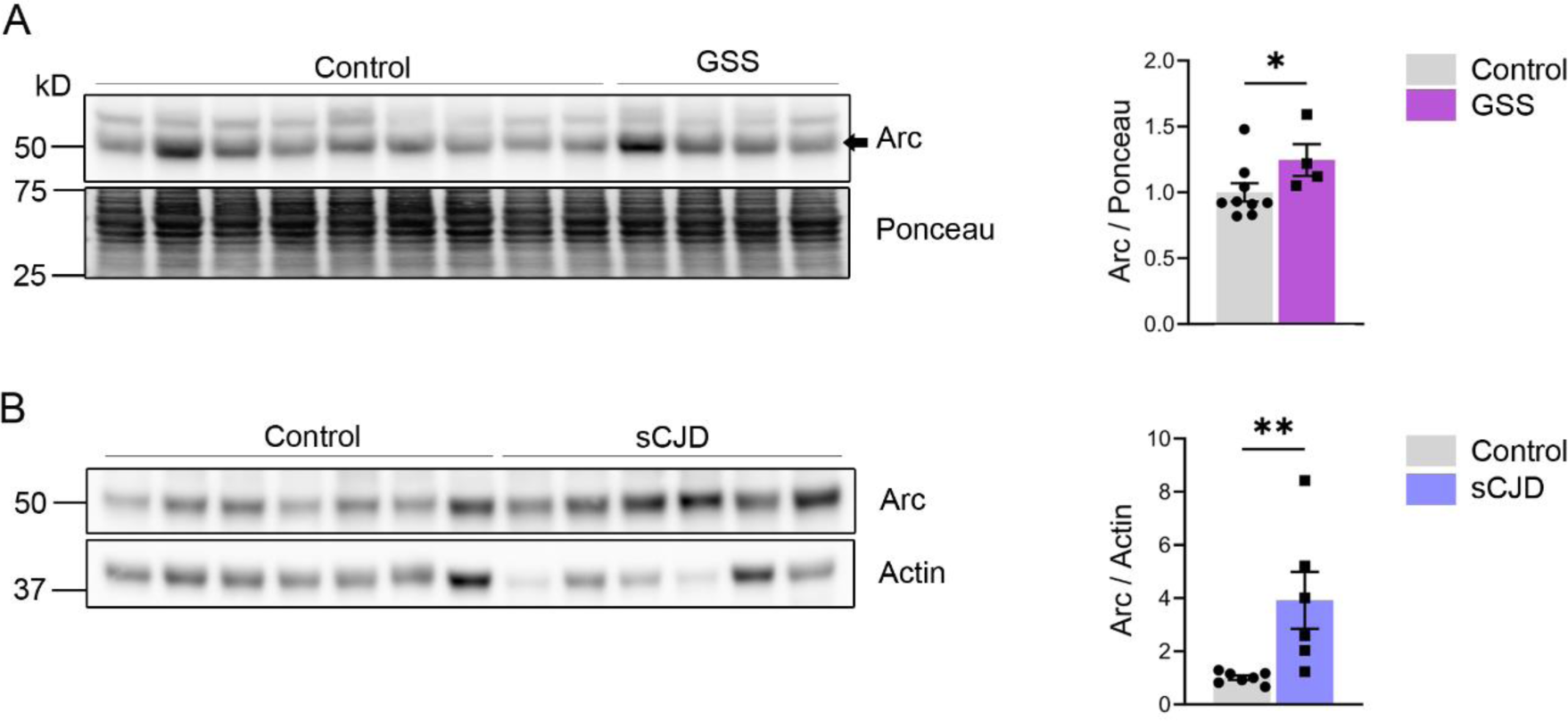
Arc is increased in human genetic and sporadic and prion disease. Immunoblotting of frontal cortex of **(A)** Gerstmann–Sträussler–Scheinker syndrome (GSS) and **(B)** sporadic Creutzfeldt-Jakob (sCJD) patients for Arc protein levels compared to age-matched controls (control: n = 7-9; sCJD: n = 6; GSS: n = 4). Data is represented as fold change compared to control samples. Mann-Whitney test. *P ≤ 0.05; **P ≤ 0.01.

**Table 1.** Number of subjects, sex, postmortem interval, and age at the time of death for control, sCJD, and GSS subjects for samples used in Figure 1. Age is shown as mean ± standard deviation (SD). Statistical analysis of age was performed using a Welch’s t-test.

| Frontal cortex sCJD |  |  |  |
| --- | --- | --- | --- |
| Characteristics | Control | sCJD | *P* value |
| Number of subjects | 7 | 6 | - |
| Age at death (years) | 72 ± 6 | 68 ± 9 | 0.3 |
| Postmortem interval (hours) | 10 ± 10 | 61 ± 41 | 0.03 |
| Sex [male (%)] | 4 (57%) | 3 (50%) | - |
| Frontal cortex GSS |  |  |  |
| Number of subjects | 8 | 4 | - |
| Age at death (years) | 73 ± 5 | 55 ± 21 | 0.2 |
| Postmortem interval (hours) | 10 ± 10 | - | - |
| Sex [male (%)] | 4 (50%) | 4 (100%) | - |

### PrP^C^ stimulation with prion oligomers or an anti-PrP^C^ antibody induces Arc in mouse cortical neurons by 2 hours

To assess whether exposure of neurons to brain-derived PrP^Sc^ impacts Arc expression, we exposed primary mouse cortical neurons to partially purified prions or mock brain for 0.5, 2, 6, 24, or 48 hours (Fig. 2A - C). By two hours post-PrP^Sc^ exposure, Arc levels increased by 2.2 – fold, then returned to baseline over the next four hours (Fig. 2A - C), indicating a rapid Arc response to PrP^Sc^.

**Figure 2.**
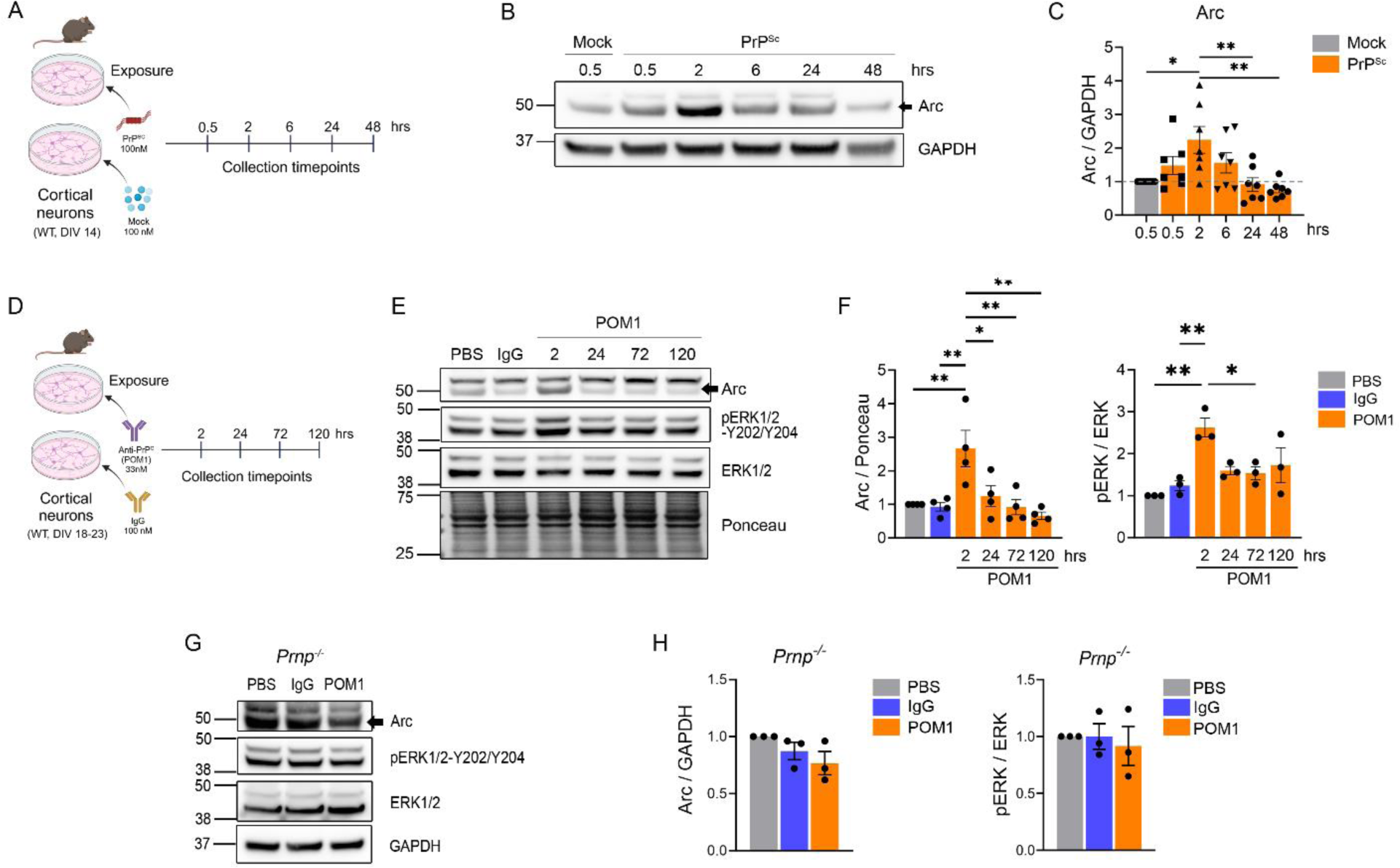
Arc is increased in response to PrP^Sc^ and anti-PrP antibody (POM1) in primary mouse cortical neurons. **(A)** Experiment schematic of PrP^Sc^ exposure of primary mouse cortical neuron culture**. (B - C)** Western blots and quantification of mouse primary cortical neurons exposed to 100 nM of PrP^Sc^ for Arc compared to mock (n = 7 independent experiments). **(D - F)** Immunoblots and quantification of mouse neurons exposed to POM1 (33 nM) (DIV 18-23; n = 4 independent experiments). **(G - H)** Immunoblots and quantification of *Prnp^-/-^* mouse cortical neurons for Arc and pERK (n = 3 independent experiments). Data is normalized to PBS for each experiment (PBS = 1). One-way ANOVA with Tukey’s MCT. *P ≤ 0.05; **P ≤ 0.01.

PrP^Sc^ partially purified from brain homogenate contains lipids and potential other molecules that bind PrP^Sc^, such as heparan sulfate. We next used a reductionist approach and tested whether triggering PrP^C^ using a well-characterized anti-PrP^C^ antibody (POM1) impacts Arc in mouse cortical neurons. POM1 binds residues 140–147 (β1 - α1 loop) and K204, R208, and Q212 on the α3 helix of the globular domain of human PrP^C^ (45) and has been shown to be a PrP^Sc^ surrogate, faithfully reproducing prion disease features including neuronal loss, protein kinase RNA-like endoplasmic reticulum kinase (PERK) activation, production of reactive oxygen species, reduced PIKfyve kinase, as well as altered PrP^Sc^-induced transcript expression in cerebellar organotypic cultured slices (COCS) (41, 42). We treated primary neurons (DIV21) for 2, 24, 72, and 120 hours with POM1 or mouse IgG [33 nM, previously reported to induce dendritic beading (46)]. Similar to PrP^Sc^ exposure, treatment with POM1 led to a 2.7 – fold increase in Arc by 2 hours (Fig. 2D - F), indicating that POM1 reproduces the Arc response following PrP^Sc^ exposure. Exposing *Prnp^-/-^* mouse cortical neurons (47) to POM1 for 2 hours failed to induce an Arc response, indicating that PrP^C^ expression was required (Fig. 2G - H; Fig. S1).

Activation of extracellular regulated kinase (ERK1/2) regulates Arc expression following neuronal stimulation by activating the nuclear kinase mitogen- and stress-activated protein kinase 1 (MSK1), which phosphorylates histone H3 at the Arc/Arg3.1 promoter (14, 48). ERK1/2 phosphorylation is increased in prion-infected mice (49). To determine if ERK1/2 activation was altered following PrP^C^ stimulation, we measured the levels of ERK1/2 phosphorylation at Thr202/Tyr204 (pERK-T202/Y204) (50). Similar to Arc, pERK1/2-T202/Y204 was significantly increased by 2 hours post-exposure to POM1 (Fig. 2D-F). ERK1/2 activation also was PrP^C^-dependent, as POM1-treated *Prnp^-/-^* neurons lacked an increase in pERK1/2-T202/Y204 (Fig. 2G - H; Fig. S1). Together these data suggest the PrP^Sc^-mimetic antibody, POM1, increases pERK1/2 and Arc in mouse cortical neurons in a PrP^C^-dependent manner.

### Stimulation of PrP^C^ in human iNs leads to an increase in Arc within 2 hours

Mouse primary neurons are a powerful tool, yet there are species-specific differences between rodent and human neurons. To define the early signaling events specific to human neurons following PrP^C^ stimulation, we used human iNs differentiated from iPSCs containing a Tet-ON 3G-controlled *Ngn2* transgene (51). To first validate the model, we measured the expression of PSD95 (postsynaptic), synaptophysin (presynaptic), and GluNB and GluN2A (glutamatergic receptors) at 3 days, 2-, 4-, and 6 - weeks post-differentiation (Fig. S2A - B). GluN2B, PSD95, and synaptophysin reached a plateau by 6-weeks or earlier (Fig. S2A - B), congruent with the appearance of synapses. Interestingly, PrP^C^ increased over time (Fig. S2A - B).

At 6 weeks, RNA-seq of the iNs showed expression markers of excitatory neurons (*SLC17A6, SLC17A7, DLG4, TUBB3*), glutamatergic receptors (*GLUN1, GRM5*), and PrP^C^ (*PRNP*) (Fig. S2C). Hitherto, the cultures had low to no expression of markers of pluripotent stem cells (*NANOG, FOXD3, POU5F1*), glial cells (*GFAP, AIF1, OLIG1, OLIG2, SOX10, MAG,* and *MOG*), or inhibitory neurons (*GAD1, GAD2*) (Fig. S2C). By immunofluorescence staining, the iNs expressed markers of mature neurons (MAP2, Tuj1, and NeuN), vesicular glutamate transporters (vGlut), and post-synaptic protein (PSD95) (Fig. S2D). PrP^C^ was expressed in the cell body and in distinct puncta around the Tuj1 positive neurites, congruent with synaptic localization of PrP^C^ (Fig. S2D).

To determine whether triggering PrP^C^ induces Arc in human neurons, we treated iNs with POM1 or mouse IgG (33 nM) from 2 to 120 hours. Strikingly, Arc was increased by 2.4 - fold after a 2-hour incubation with POM1, while PrP^C^ was reduced, indicating that an Arc response could be acutely triggered by a PrP^C^ ligand (Fig. 3; Fig. S3). pERK1/2-Thr202/Tyr204 was also increased (Fig. 3; Fig. S3).

**Figure 3.**
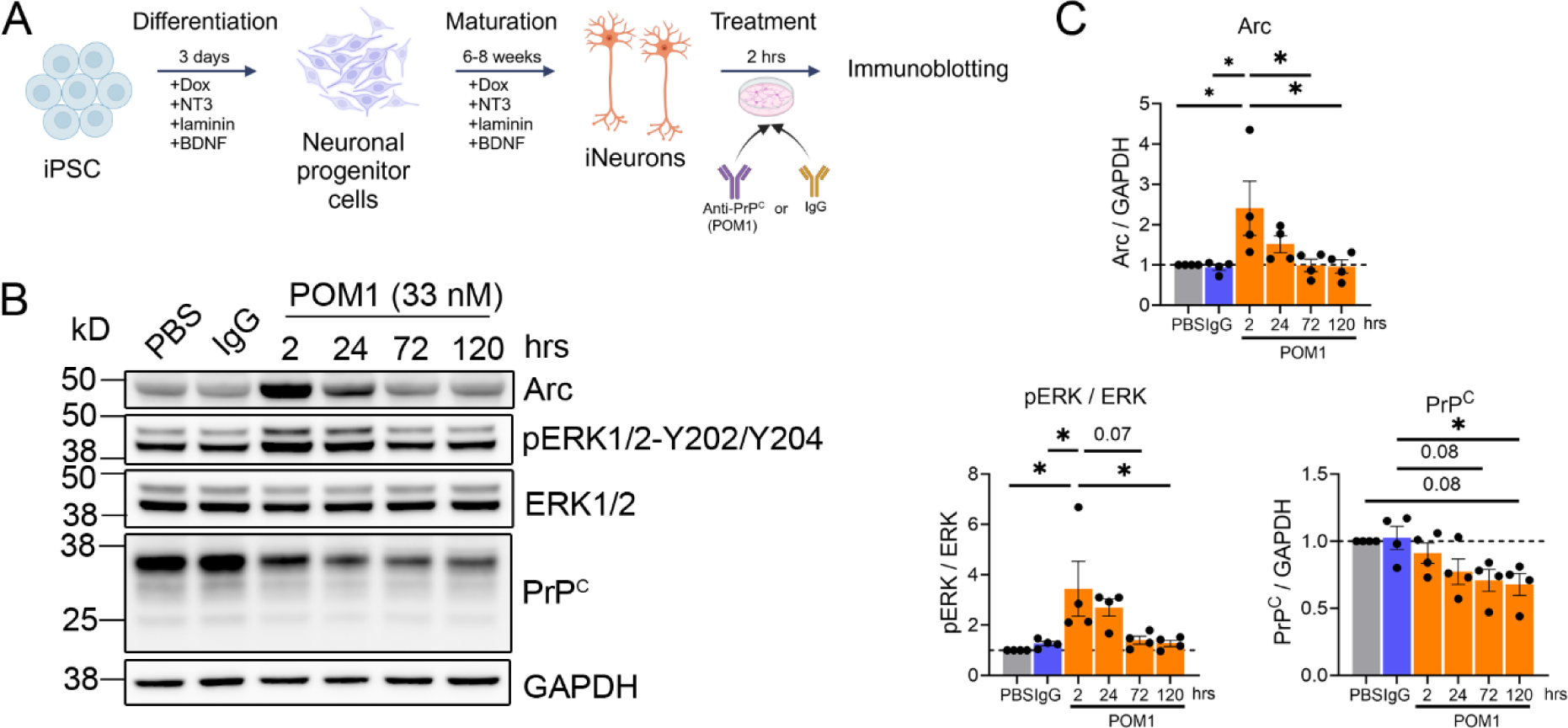
Human iNs show early ERK activation and increased Arc following POM1 stimulation. **(A)** Schematic timeline of differentiation, maturation, and exposure to POM1. **(B - C)** Western blot and quantification of iNs stimulated with POM1 (33 nM) (n = 4 independent experiments). Panel C, data is normalized to PBS for each experiment (PBS = 1), one-way ANOVA with Tukey’s MCT. *P ≤ 0.05; **P ≤ 0.01.

### Stimulation of PrP^C^ alters EGFR signaling by 2 hours

To identify membrane receptors and kinases that may underlie the increase in Arc in an unbiased manner, we employed an array approach. We exposed iNs to POM1 or an IgG control for 2 hours, applied the lysate to a membrane-based antibody array, and measured the relative levels of 37 phospho-kinase phosphorylated proteins. We found a significant increase in phosphorylated phospholipase C (pPLC)-γ1-Y783, and a decrease in phosphorylated (p) EGFR-Y1086 (a tyrosine kinase), as well as pAKT-T308, p70 S6 kinase-T389, peNOS-S1177, pp53-S15/S46/S392, pp38-T180/Y182, pSrc-Y419, pPYK2-Y402 (Fig. 4A - B; Fig. S4), supporting that POM1-triggered stimulation of PrP^C^ triggers rapid intracellular signaling.

**Figure 4.**
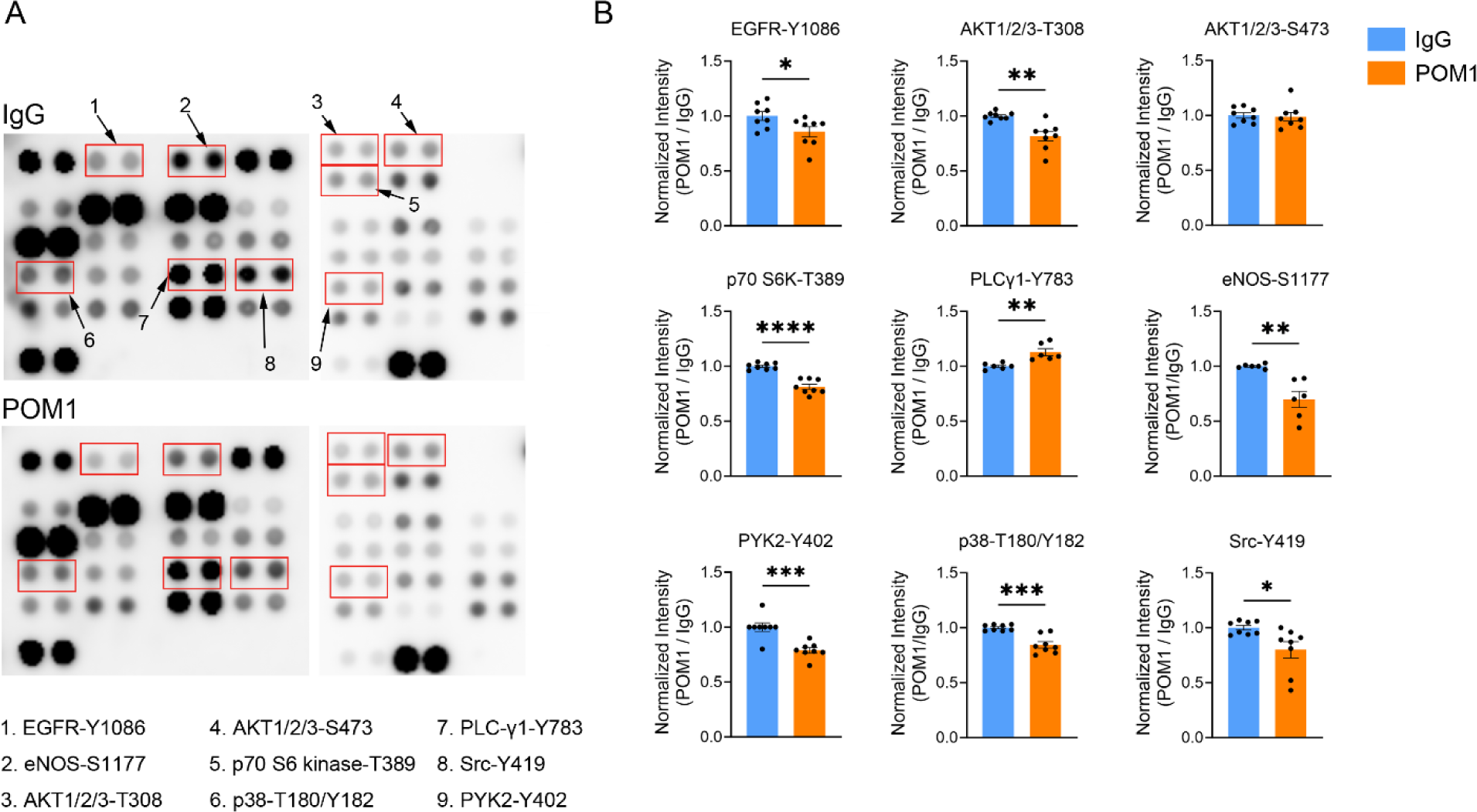
Protein phosphorylation in POM1-treated iNs. **(A)** Phospho-kinase array of 37 kinases and related proteins in 6 - 8 week-old iNs treated with POM1 or IgG for 2 hours. **(B)** Quantification of (A) (n = 4 independent experiments, each with 2 technical replicates). Data is normalized to the average of the two technical replicates of IgG for each experiment, Welch’s t-test. *P ≤ 0.05; **P ≤ 0.01.

### Prion mimetic antibody triggers transcriptional changes associated with prion disease and EGFR signaling

To identify RNA and protein alterations occurring in parallel with the increase in Arc, pERK1/2, and pPLC-γ1, we performed RNA sequencing (RNA-seq) and proteomics of iNs exposed to POM1 or IgG for 2 hrs. The RNA-seq analysis revealed 53 differentially expressed genes (DEGs) (increased = 29, decreased = 24; p ≤ 0.1; ± 25% fold-change) (Fig. 5A - B; Table S1). Of those 53 genes, 15% were long non-coding RNAs and 17% were involved in EGFR signaling (Fig. 5A – B; Table S1). The downregulated EGFR-related genes include *CGRRF1*, an E3-ubiquitin ligase necessary for the proteasomal degradation of EGFR (53), as well as *KLF11,* a transcriptional repressor negatively regulated following EGFR activation (54, 55). The upregulated EGFR-related transcripts included modulators of the EGFR ligand (EGF) (*GRPR* and *PTGER3*) and EGFR transcription (*TFAP2C*), as well as mediators of EGFR signaling (*DNMT3A*, *PTPN11*, *RASGRP1*, and *DGKI*) (Fig. 5A - B; Table S1). These DEGs implicate altered EGFR signaling as a cellular response to PrP^C^.

**Figure 5.**
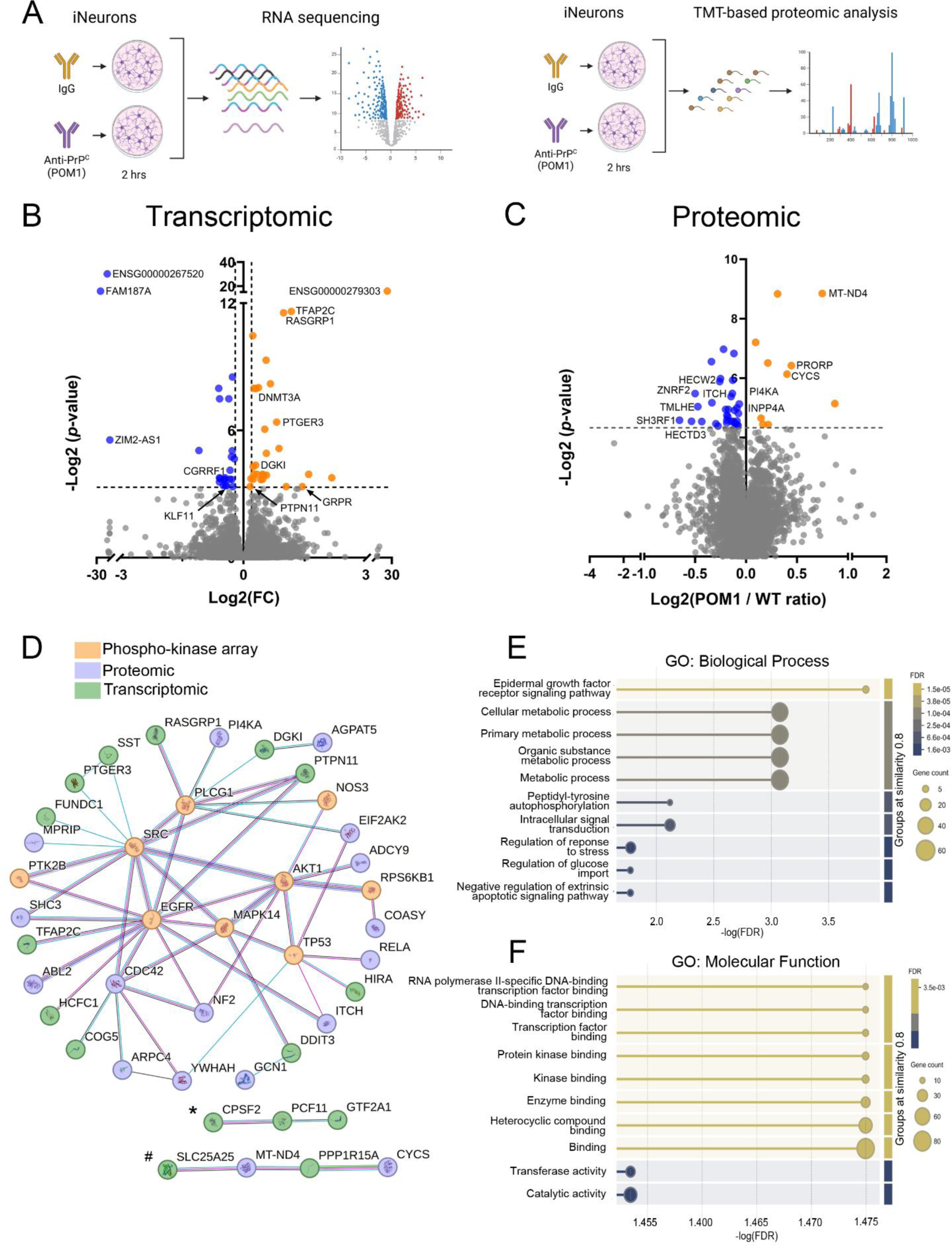
Multiomic profiling of iNs following exposure to POM1. **(A)** Schematic representation of RNA-sequencing and of TMT-based proteomic analysis of iN treated with POM1 or IgG for 2 hours. **(B)** Volcano plot shows genes significantly altered following POM1 or IgG exposure (n = 4 independent experiments; blue = significantly decreased; orange = significantly increased). **(C)** Volcano plot showing significantly altered peptides following POM1 or IgG treatment (n = 4 independent experiments; blue = significantly decreased; orange = significantly increased). **(D)** Predicted network of protein-protein interactions of significantly altered genes and proteins from our transcriptomic (green), proteomic (blue), and phospho-kinase array analysis (orange). Three clusters are observed. EGFR is the central node (most predicted interactions) of network 1. * and # depict networks related to transcription factor and mitochondrial proteins, respectively. **(E - F)** Biological Process and Molecular Function Gene Ontology (GO) terms from protein-protein interaction analysis.

The RNA-sequencing also revealed 12 DEGs (23%) that were altered in POM1-treated iNs that were reported to be altered in the hippocampus of prion-infected mice (Fig. 5A - B; Table S1). These include *SST*, *LSM11*, *AATK*, *ABCD2*, *SATB2*, and *GRPR* in prion-infected (RML stain) hippocampus, *RNF138*, *HCFC1*, and *FUNDC1* in prion-infected (RML stain) hippocampus (CA1 only), and *SLC25A25* in prion-infected hippocampus (22L stain) (10, 56, 57). Neurotoxic factor PTGER3, which contributes to prion disease pathology and regulates EGFR ligands, was also increased (58).

### Triggering PrP^C^ induces proteomic alterations linked to metabolism

To further probe how triggering PrP^C^ acutely impacts downstream signaling, we performed tandem mass tag (TMT) - labeled mass spectrometry on iNs following exposure to POM1 or IgG for 2 hours. We detected 4391 proteins, with 43 being differentially expressed (n = 10 increased, n = 33 decreased) (p ≤ 0.05) (Fig 5A, C; Table S2). Notably, there was an increase in mitochondrial proteins (MT-ND4, CYCS, and PRORP) and a decrease in ubiquitin E3-ligases (HECW2, ZNRF2, ITCH, HECTD3, and POSH), suggesting that triggering PrP^C^ impacts metabolic and protein degradation pathways.

There was also a decrease in phosphatidylinositol 4-kinase (PI4K) and inositol polyphosphate-4-phosphatase type I A (INPP4A) peptides. PI4K and INPP4A are components of the phosphoinositide pathway (Fig. 5A, C) and a reduction would be expected to reduce the synthesis of secondary messengers, such as PI4P and PtdIns(3,4)P2 (59, 60), suggesting that PrP^C^ stimulation impacts metabolic activity.

### Network analysis reveals three protein-protein interaction clusters

To visualize the possible protein-protein interactions of our significantly altered phospho-kinase, transcriptional, and proteomic signatures following PrP^C^ stimulation, we used StringDB database. This analysis identified three clusters: (i) an EGFR signaling network (with EGFR as the central node), (ii) a network of mitochondrial proteins, and (iii) a network of transcription factors and mRNA processing (Fig 5D). Gene Ontology (GO) enrichment analysis of the Biological Process pathways revealed “Epidermal growth factor receptor signaling pathway” as the top altered pathway (False Discovery Rate = 0.00015; Fig. 5E). GO analysis of Molecular Function pathways revealed “protein kinase binding” as significantly enriched (Fig. 5F).

### Phospho-PLC-γ1 and phospho-EGFR are altered in late stages of prion disease

To test whether pPLC-γ1 and EGFR signaling are altered in vivo, we inoculated two cohorts of WT mice with either 22L or RML prion strains, or an uninfected (mock) brain control, and collected and analyzed cerebral cortex at the start of clinical signs of prion disease (22L: 80% or RML: 70% of disease progression) by western blot. We observed a significant increase in pPLC-γ1-Y783 (56 ± 15.6% and 40 ± 14.1% increase in 22L- and RML-prion-infected mice, respectively) compared to the mock-inoculated controls (Fig. 6A - D). There was a significant decrease in pEGFR-Y1086 (52 ± 4.8% and 35 ± 8.9% reduction in 22L- and RML-prion-infected cortex, respectively). These results suggest that prion infection increases PLC-γ1 activity in the brain *in vivo*, while reducing EGFR phosphorylation, and shows the predictive power of the POM1-iN model for studying PrP-induced signaling. Taken together, these data suggest a model in which triggering PrP^C^ with POM1 or prions increases active PLC-γ1 in human neurons and prion-infected brain, respectively, which would cleave PIP2 into IP3. IP3 releases Ca^2+^ from ER stores, and high cytosolic Ca^2+^ induces an Arc response to reduce accessible AMPAR at the synaptic membrane and lower neuronal activity.

**Figure 6.**
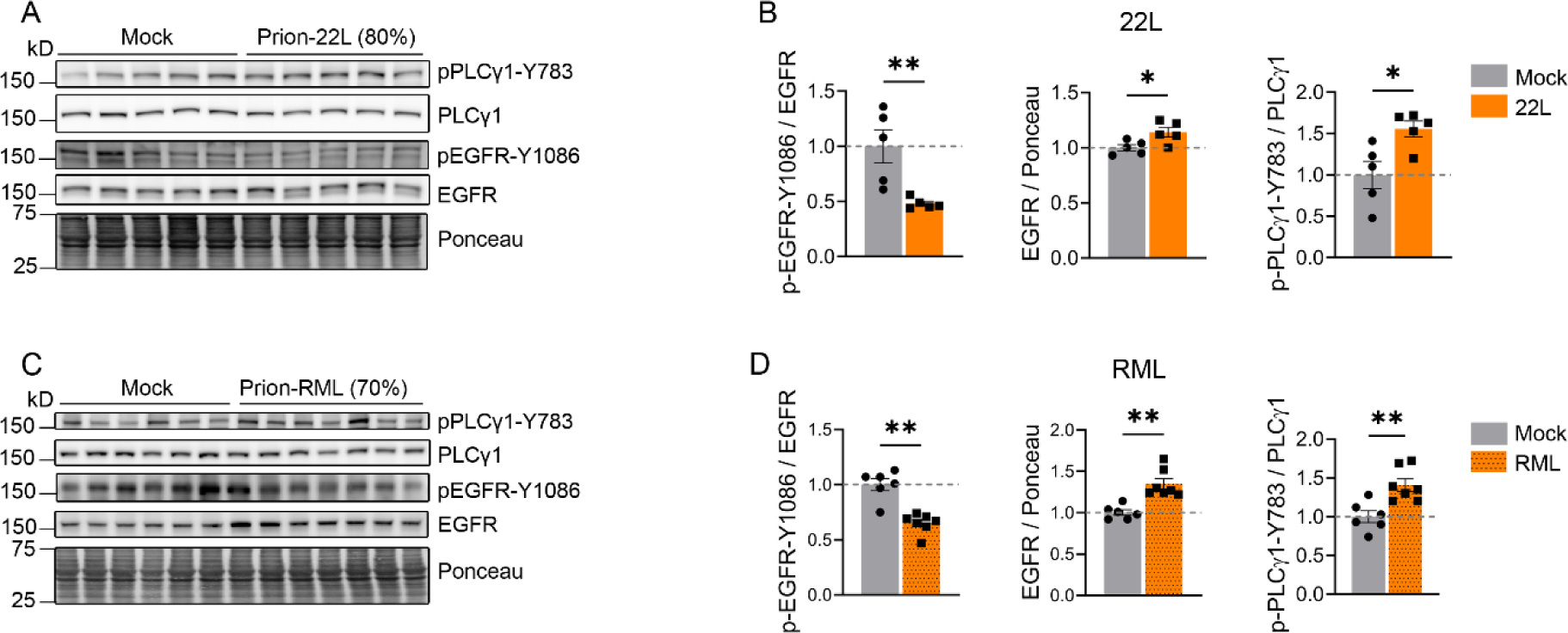
EGFR signaling is altered in prion-affected brains. **(A - D)** Western blotting and quantification of prion-infected mouse brain collected at the onset of clinical signs. Cortices from two prion strains (22L and RML) were compared to mock-inoculated controls (mock: n = 5 - 6; 22L: n = 5; RML: n = 7). Data is represented as fold change compared to mock samples. One-way ANOVA with Tukey’s MCT. *P ≤ 0.05; **P ≤ 0.01

## Discussion

GPI-anchored proteins commonly localize to lipid rafts and interact with cell surface receptors to signal intracellularly (61). Here we show that a pathogenic PrP^C^ ligand in human iNs triggers an intracellular signaling response, increasing PLC-γ1 phosphorylation, an upstream regulator of endoplasmic reticulum (ER) Ca^2+^ release (62), and expression of the activity response gene, *Arc*. We also provide the first evidence that PLC-γ1 phosphorylation is increased in the brain of prion-infected mice at the start of clinical symptoms.

POM1-induced stimulation of PrP^C^ induced PLC-γ1 phosphorylation in human iNs, within two hours. PLC-γ1 activation catalyzes the production of inositol-1,4,5-trisphosphate (IP3), which releases ER Ca^2+^ stores (62). While PLC-γ1 activation is not directly known to regulate Arc, its phosphorylation and the subsequent increase in intracellular Ca^2+^ could promote an increase in Arc (52). The resulting increase in Arc may serve as a homeostatic response, regulating synaptic strength by promoting GluA1 internalization in weakly active synapses (16, 63).

We reveal an increase in phosphorylated PLC-γ1 in prion-infected mice at the onset of clinical symptoms. Chronic activation of PLC-γ1 could result in a prolonged increase in cytosolic Ca^2+^ (64), which has been linked to neurotoxicity (49, 65, 66). For example, excessive ER Ca^2+^ release has been implicated in NMDA-driven excitotoxicity (66). Further research is needed to elucidate the disease stage and consequences of sustained PLC-γ1–PIP2 cleavage and prolonged high intracellular Ca^2+^ in prion disease.

We used human iNs as a human-relevant model to study early signaling events following PrP^C^ stimulation, recognizing that there are limitations. We reveal an increase in PLC-γ1 and Arc, however, the subcellular localization remains to be elucidated. Additionally, while our study examined acute PrP^C^ stimulation, prion diseases develop over years. The long-term impact of chronic PLC-γ1 signaling alterations remains unclear and requires further investigation. Furthermore, while Arc serves as a well-established marker of neuronal activity (14, 15), additional functional assays measuring synaptic plasticity and network excitability would strengthen our understanding of neuronal hyperactivity and dysfunction in prion disease.

Using an antibody that measures activation of EGFR based on phosphorylation of Tyr-1086, the Grb2-binding site (67), we showed that in iNs, POM1 decreases activation of the EGFR. Similarly, EGFR Tyr-1086 was decreased in late-stage prion-infected mouse brains. Groveman et al (68) showed decreased EGFR in PrP^C^-knock-out neural stem cells and linked this correlation to increased cellular senescence *in vitro*. Thus, interactions between the EGFR and PrP^C^ derivatives may impact normal physiology and prion disease progression. Factors that may regulate the PrP^C^ - EGFR interaction include gene transcription, trafficking of the EGFR to the cell surface and between membrane microdomains, and diverse pathways that regulate the availability of EGFR ligand. Understanding this interaction is an important future goal.

In summary, we show that Arc, a marker of neuronal activity, is elevated in both sporadic and familial human prion disease, and rapidly increases in human iNs following PrP^C^ stimulation. While further studies are necessary to evaluate the role of PLC-γ1, our findings suggest that triggering PrP^C^ rapidly impacts intracellular signaling leading to increased neuronal activity markers.

## Methods

### Prion inoculation of mice

C57BL/6J mice (6 – 8 weeks old) were anesthetized with ketamine and xylazine and inoculated into the left parietal cortex with 30 μl of 1% 22L or RML prion-infected brain homogenate prepared from terminally ill mice or 1% mock brain homogenate prepared from uninfected mice. The 22L and RML strains are mouse-adapted prions originally derived from sheep scrapie and characterized by diffuse prion aggregates, synaptic and neuronal loss, and glial activation in the brain (69–72). Mice were maintained under pathogen-free conditions on a 12:12 light/dark cycle, and with access to standard laboratory chow and water ad libitum.

Prion-infected or mock-inoculated mice (n = 5-7) were euthanized at the beginning of clinical signs (22L: 80% of disease progression; 120 d.p.i and RML: 70% of disease progression; 123 d.p.i). The timepoint were approximated based on previous studies using each strain, however, the actual time to terminal disease may differ among cohorts. The cortex was dissected and flash-frozen in liquid nitrogen for immunoblotting and biochemical experiments.

### Immunoblotting of mouse cortex

Cortices were homogenized in RIPA buffer (10% w/v) using a Beadbeater™ tissue homogenizer. Samples were lysed on ice for 30 min using 2% N-lauryl sarcosine in PBS with Phos-STOP™ (Pierce), nucleases [Benzonase™ (Sigma)], and protease inhibitors (Complete TM), then centrifuged at 2000g at 4 °C for 10 min. Proteins in the supernatant were quantified by bicinchoninic acid (BCA) assay, and equivalent amounts of protein were electrophoresed through a 10% Bis-Tris gel (Invitrogen) and transferred to a nitrocellulose membrane by wet blotting. Membranes were incubated with primary antibodies overnight at 4 °C followed by incubation with an HRP-conjugated secondary antibody. The immunoblots were developed using a chemiluminescent substrate (ECL Supersignal West Dura or Femto, ThermoFisher Scientific) and visualized on a Fuji LAS 4000 imager. Densitometry analysis was performed using MultiGauge software (Fujifilm).

### Patients with sporadic Creutzfeldt-Jakob disease and controls

The sCJD brain tissues used in this study were obtained from a prospective study of sCJD patients evaluated between July 2015 and 2018 at the University of California, San Francisco (UCSF) Memory and Aging Center clinical research program for rapidly progressive neurologic disease (Table 1). All patients with sCJD had extensive clinical testing, including brain MRI, CSF analysis for 14-3-3, neuron-specific enolase (NSE), and total tau, and were classified premortem as probable sCJD by UCSF clinical and radiologic diagnostic criteria (Geschwind et al., 2007; Staffaroni et al., 2017). Cases were diagnosed postmortem as sCJD by Western blotting. Autopsy-verified, nonpathologic, aged (control) brain samples were obtained from the Neuropathology Core and brain bank of the Shiley-Marcos Alzheimer’s Disease Research Center at the University of California, San Diego (UCSD; Table 1). These control brains were devoid of significant neurodegenerative disease pathology (postmortem) and disease-related cognitive impairment at the antemortem assessment proximate to death. This study was approved by the ethics committee at UCSF. Informed consent was received by all patients (IRB Study number 10-04905). All brain tissues used were de-identified samples collected at autopsy (NIH Office of Human Subjects Research Protections, exemption 4).

### Immunoblot analysis of sCJD and GSS brain samples

Immunoblotting of frontal cortex from sCJD and GSS patients was performed as previously published with minor modifications (10). Briefly, frontal cortex was homogenized in PBS using a Beadbeater tissue homogenizer. Protein levels were quantified using a BCA protein assay, 30 µg of protein per sample was lysed in PBS containing 2% sarcosyl, endonuclease (benzonase), phosphatase inhibitors (PhosStop), and protease inhibitors (Complete Mini) for 15 min at 37°C, and then centrifuged for 30 s at 18,000 × g to remove debris. Samples were boiled for 5 min in LDS loading buffer (Invitrogen) and electrophoresed through a 10% Bis-Tris gel (Invitrogen) prior to transfer to a nitrocellulose membrane.

### Induced-pluripotent stem cell culture and neuronal differentiation

Human iPSC from the WTC11 line containing a Tet-ON 3G-controlled *Ngn2* transgene (51) were cultured in DMEM/F-12 (HAM) with Essential 8 supplement. iNs were differentiated with a two-step protocol (pre-differentiation and maturation) as previously described by (51). Briefly, for pre-differentiation, iNs were plated in six-well plates coated with Matrigel, and incubated in knockout Dulbecco’s modified Eagle’s medium (KO-DMEM)/F12 medium containing N2 supplement, non-essential amino acids (1X NEAA; Life Technologies™), mouse laminin (1 μg/mL; Life Technologies™), brain-derived neurotrophic factor (BDNF, 10 ng/mL; Peprotech), neurotrophin-3 (NT3,10 ng/mL; Peprotech), doxycycline (2 μg/mL; Sigma) for 3 days and Y-27632 (10 µM; Cayman). The medium was changed daily, and Y-27632 was removed from day 2. For maturation, pre-differentiated iN precursor cells were dissociated, counted, and plated at 650,000 cells/well in six-well plates coated with poly-L-lysine (PLL) in maturation medium containing 50% DMEM/F12, 50% Neurobasal-A medium, 0.5X B27 supplement, 0.5X N2 supplement, 1X GlutaMax, 1X NEAA, doxycycline (2 μg/mL), mouse laminin (1 μg/mL), BDNF (10 ng/mL), and NT3 (10 ng/mL). Half the medium was changed weekly.

### iN treatment

After 6-8 weeks of maturation, the iNs were treated with the anti-PrP antibody POM1 or IgG (33 nM) for the indicated time points.

### Primary neuron culture

Primary cortical neurons were isolated from postnatal day (P)0-1 C57BL/6J or *Prnp^-/-^* mice pups. The cerebral cortices were dissected in dissection buffer (1 m MgCl2, 1 m CaCl2,1 M HEPES, and 2.77 M glucose), dissociated with 0.25% trypsin at 37°C for 20 min, treated with DNase, and triturated. Debris was removed by passing the cells through a 70-µm cell strainer. Cells were then centrifuged for 10 min and cultured in neurobasal media containing 2% B27-Plus supplement and 1X GlutaMAX.

C57BL/6J or *Prnp^-/-^* cell at day *in vitro* (DIV) 21 cells were treated with POM1 or IgG (33 nM). C57BL/6J cells at DIV 14 cells were treated with partially purified 22L or mock brain homogenate (100 nM) for the indicated time. Cells were collected on ice using lysis buffer (10 mM Tris-HCl, 150 mM NaCl, 10 mM EDTA, 0.5% NP40, 0.5% DOC; pH 7.4 with Phos-STOP™, nucleases [Benzonase™], and protease inhibitors (Complete™). Lysates were cleared by centrifugation at 10,000 x g for 10 min at 4 °C. Proteins in the supernatant were quantified by BCA assay, and equivalent amounts of protein were electrophoresed through a 10% Bis-Tris gel.

### Partially purified of prion oligomers

Prion-infected and uninfected 10% brain homogenates in PBS were lysed by adding an equal volume of 2% N-lauryl sarcosine in PBS supplemented with benzonase and 1 mM MgCl2 (final concentration). Lysates were incubated for 30 min at 37 °C with mixing, and centrifuged at 18,000 g for 60 min at 4°C. The pellets were carefully rinsed with PBS, centrifuged at 18,000 g for 10 min at 4°C, re-suspended in PBS, and heated to 65 °C in preparation for cell culture. The PrP concentration was measured for each preparation by quantifying levels via immunoblotting, using a standard curve from a serial dilution of recombinant PrP.

### RNAseq

For RNAseq, 6 - 8 week-old iNs were treated for 2 hrs with PBS, POM1 (33 nM), or IgG (33 nM). Samples were lysed on RLT buffer and isolated RNeasy Micro kit (Cat# 74004; Qiagen) following the manufacture’s protocol. Total RNA was assessed for quality using an Agilent Tapestation 4200, and samples with an RNA Integrity Number (RIN) greater than 8.5 were used to generate RNA sequencing libraries using the TruSeq Stranded mRNA Sample Prep Kit (Illumina, San Diego, CA). Samples were processed following the manufacturer’s instructions. Resulting libraries were multiplexed and sequenced with 100 base pair (bp) single reads (SR100) to a depth of approximately 25 million reads per sample on an Illumina NovaSeq X Plus. Samples were demultiplexed using bcl2fastq Conversion Software (Illumina, San Diego, CA). QC and RNAseq were conducted at the IGM Genomics Center, University of California, San Diego, La Jolla, CA.

### Phosphoprotein array studies

Phosphoproteins were detected using the Proteome Profiler™ human phospho-kinase array (R&D systems Cat# ARY003C) following the manufacturer’s instructions with minor modifications. Briefly, 6 - 8 week-old iNs treated for 2 hrs with POM1 or IgG (33 nM) were lysed using kit lysis buffer (lysis buffer 6) with protease inhibitors (Complete™). Membranes were incubated with 600 µg of protein overnight at 4 °C, followed by incubation with detection antibody cocktails, and streptavidin-HRP for detection. The immunoblots were developed using a chemiluminescent substrate (ECL Supersignal West Dura, ThermoFisher Scientific) and visualized on a Fuji LAS 4000 imager. Densitometry analysis was performed using MultiGauge software (Fujifilm).

### RNA-seq data processing

RNA-sequencing reads were trimmed of adaptor sequences using cutadapt (v1.4.0) and mapped to repetitive elements (RepBase v18.04) using the STAR (v2.4.0i). Reads that did not map to repetitive elements were then mapped to the mouse genome (mm9). GENCODE (v19) gene annotations and featureCounts (v.1.5.0) were used to create read count matrices. Differential expression analysis was performed with DESeq2 v1.32.0 on the raw counts matrix.

### Mass spectrometry

6-week old iNs were treated with POM1 or IgG for 2 hrs, then collected on ice using lysis buffer (10 mM Tris-HCl, 150 mM NaCl, 10 mM EDTA, 0.5% NP40, 0.5% DOC; pH 7.4 with Phos-STOP™, nucleases [Benzonase™], and protease inhibitors (Complete™). Lysates were cleared by centrifugation at 10,000 x g for 10 min at 4 °C. Proteins in the supernatant were quantified by BCA assay. 200 µg of samples were precipitated with methanol and chloroform, then digested with trypsin as previously described (73). The digested peptides were desalted, and then dried in a speed-vac. Each peptide sample was labeled with a unique TMT 10-plex isobaric label (Thermo Scientific) according to a published method (74).

After the TMT-labeled samples were combined into one tube, 50 µg were removed for unmodified analysis and the remaining was used for phosphorylation enrichment. Phosphopeptides were enriched using sequential titanium dioxide (TiO2) and ferric nitrilotriacetate (Fe-NTA) using a phosphorylation enrichment kit (Thermo Scientific). The two phosphorylation enrichments were combined into one sample.

The TMT unmodified and phosphorylated peptides were then each fractionated offline by high pH reverse-phase spin columns (Thermo Scientific). The TMT-labeled samples were analyzed on an Orbitrap Fusion Eclipse Tribrid mass spectrometer (Thermo Scientific). Samples were injected directly onto a 25 cm, 100 μm ID column packed with BEH 1.7 μm C18 resin (Waters). Samples were separated at a flow rate of 300 nL/min on an EasynLC 1200 (Thermo). Buffer A and B were 0.1% formic acid in water and 90% acetonitrile, respectively. A gradient of 1–25% B over 120 min, an increase to 40% B over 40 min, an increase to 100% B over 10 min, and held at 100% B for 10 min was used, for a 180 min total run time. Peptides were eluted directly from the tip of the column and nanosprayed directly into the mass spectrometer by application of 2.5 kV voltage at the back of the column. The Eclipse was operated in a data dependent mode. Full MS1 scans were collected in the Orbitrap at 120k resolution. The cycle time was set to 3 s, and within this 3 s the most abundant ions per scan were selected for CID MS/MS in the ion trap. MS3 analysis with multinotch isolation (SPS3) was utilized for detection of TMT reporter ions at 60k resolution. Monoisotopic precursor selection was enabled and dynamic exclusion was used with an exclusion duration of 10 s.

Proteome Discoverer 2.5 was used to search the MS/MS data to identify peptides, quantify the data, and perform statistical analysis. The MS spectra was searched using the Uniprot human protein database with isoforms (downloaded on 2023-2-16) and a common contaminant proteins list. The decoy database was the reverse of this Uniprot database to filter identifications to a 1% FDR. The peptides were allowed to have a maximum of two miscleavages. The static modification searched were TMT on lysine and peptide N-terminal (+229.162932 Da) and cysteine carbamidomethylation (+57.021464 Da). Phosphorylation (79.9663) was searched as a differential modification on serine, threonine, and tyrosine. Reporter ion distributions specific to the lot number of the TMT reagent were employed as correction factors. Mass spectrum raw files will be uploaded to ProteomExchange upon publication.

### Primary antibodies for western blots

The following antibodies were used for western blotting: mouse anti-PrP [1:5,000, POM2; POM1 (75)]; anti-GAPDH-HRP (1:5000, GeneTex; #GTX627408-01); anti-phosphorylated EGFR-Y1068 (1:500, Cell Signaling Technology; #3777); anti-EGFR (1:500, Cell Signaling Technology; #54359); anti-phosphorylated ERK1/2-T202/Y204 (1:5000, Cell Signaling Technology; #4370); anti-ERK1/2 (1:5000, Cell Signaling Technology; #4695); anti-phosphorylated PLCγ-1-Y783 (1:5000, Cell Signaling Technology; #14008); anti-PLCγ-1 (1:2000, Cell Signaling Technology; #5690); anti-ARC/ARG3.1 (1:5000, Proteintech, #16290-1-AP).

### Approval of animal studies

All animal studies were approved by the Institutional Animal Care and Use Committee at UC San Diego. Protocols were performed in accordance with good animal practices, as described in the Guide for the Use and Care of Laboratory Animals published by the National Institutes of Health.

### Statistical analysis

Analysis of western blotting data was performed using Prism software (GraphPad Software, Inc.). Comparisons between two groups were made by Welch’s t-test, whereas multiple groups were compared by one-way analysis of variance (ANOVA) followed by Tukey HSD post hoc test. P-values ≤ 0.05 were considered statistically significant.

## Author contributions

CJS, DOJ, and SLG conceptualized and designed experiments. DOJ, GF, ER, AJR, and KS performed experiments and analyzed the data. DBM performed mass spectrometry experiments. CJS, DOJ, SLG, XC, and JRY contributed to method design, data analysis, and interpretation. DOJ and CJS wrote the manuscript.

## Acknowledgments

We thank the National Prion Disease Pathology Surveillance Center (NPDPSC) for prion typing and all patients and their families for participating in the research. Microscopy and image analysis were performed at the Nikon Imaging Center at UC San Diego, RNAseq data was generated at the UC San Diego IGM Genomics Center. We thank Drs. Peng Guo and Richard Sánchez for the support on microscopy experiments, Dr. Kristen Jepsen for support on RNAseq experiments, the animal care staff at UC San Diego for excellent animal care, Dr. Michael Geschwind, Jeff Metcalf and the laboratory of Dr. Robert Rissman for providing patient samples.

---

**Figure S1.**
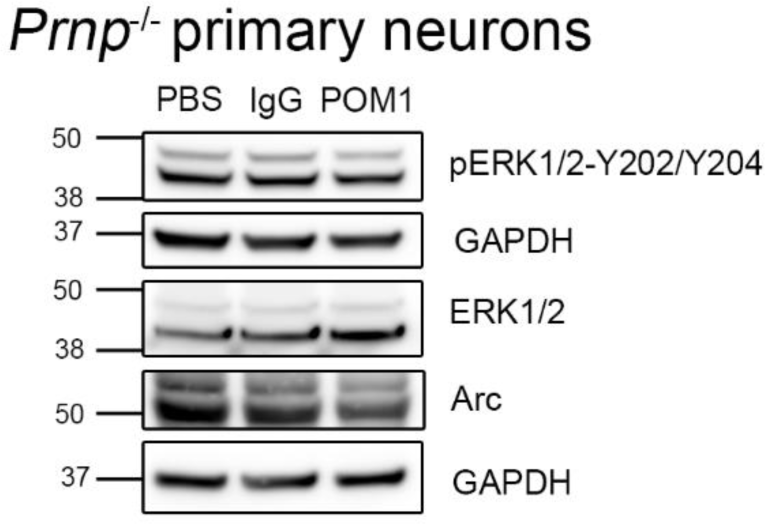
Immunoblotting of primary *Prnp^-/-^* neurons shown in Figure 1, with the corresponding GAPDH for each blot.

**Figure S2.**
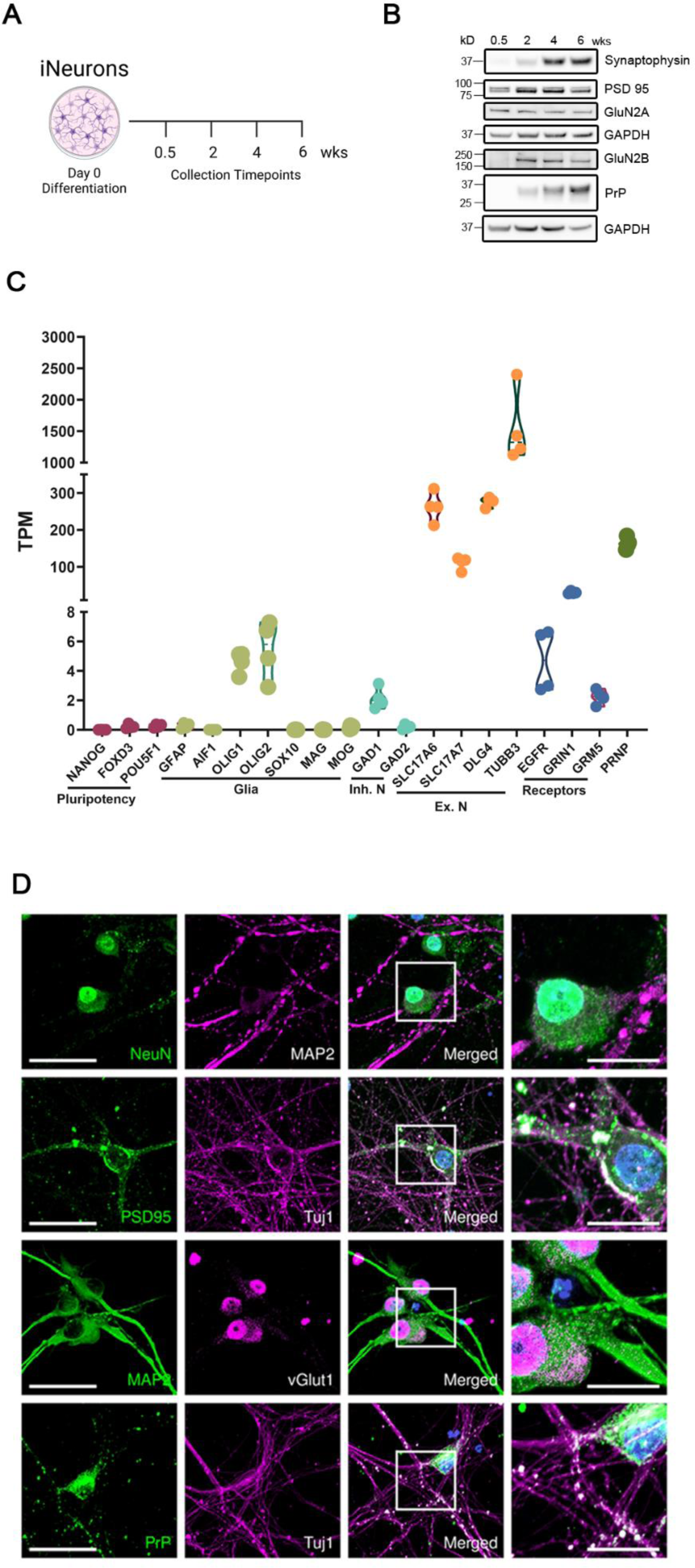
Characterization of neuronal and synaptic protein expression in human iNs. (**A - B)** Immunoblotting of synaptic proteins (postsynaptic: PSD95, presynaptic: synaptophysin), glutamatergic receptor markers (GluNB and GluN2A), and PrP^C^ in iNs throughout maturation. **(B)** RNA-seq analysis of 6-week-old iNs demonstrates expression of excitatory (*SLC17A6*, *SLC17A7*, *DLG4*, *TUBB3*) and glutamatergic (*GRIN1*, *GRM5*) markers along with the cellular prion protein coding gene (*PRNP*). N = 4 independent experiments. **(C)** Representative immunofluorescent images of 6-week-old iNs showing expression of mature neuronal markers (MAP2, Tuj1, and NeuN), vesicular glutamate transporters (vGlut), and post-synaptic protein (PSD95) along with PrP^C^ localized to the synapse. Scale bar = 50 µm. The higher magnification column (right panels) have a scale bar of 20 µm.

**Figure S3.**
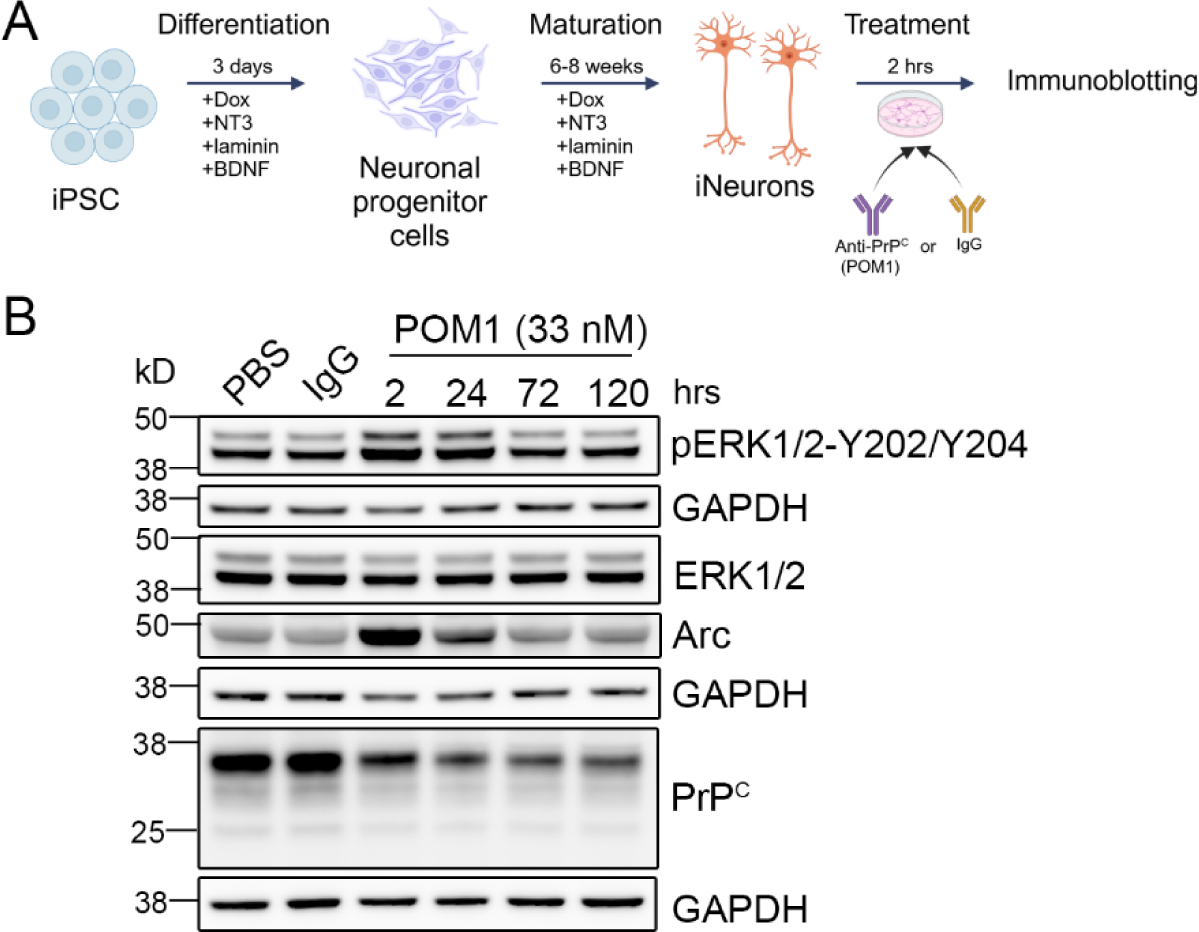
ERK activation and increase in Arc upon POM1 treatment, all loading controls (GAPDH) included. Immunoblotting of iNs shown in Figure 2, with the corresponding GAPDH for each individual blot.

**Figure S4.**
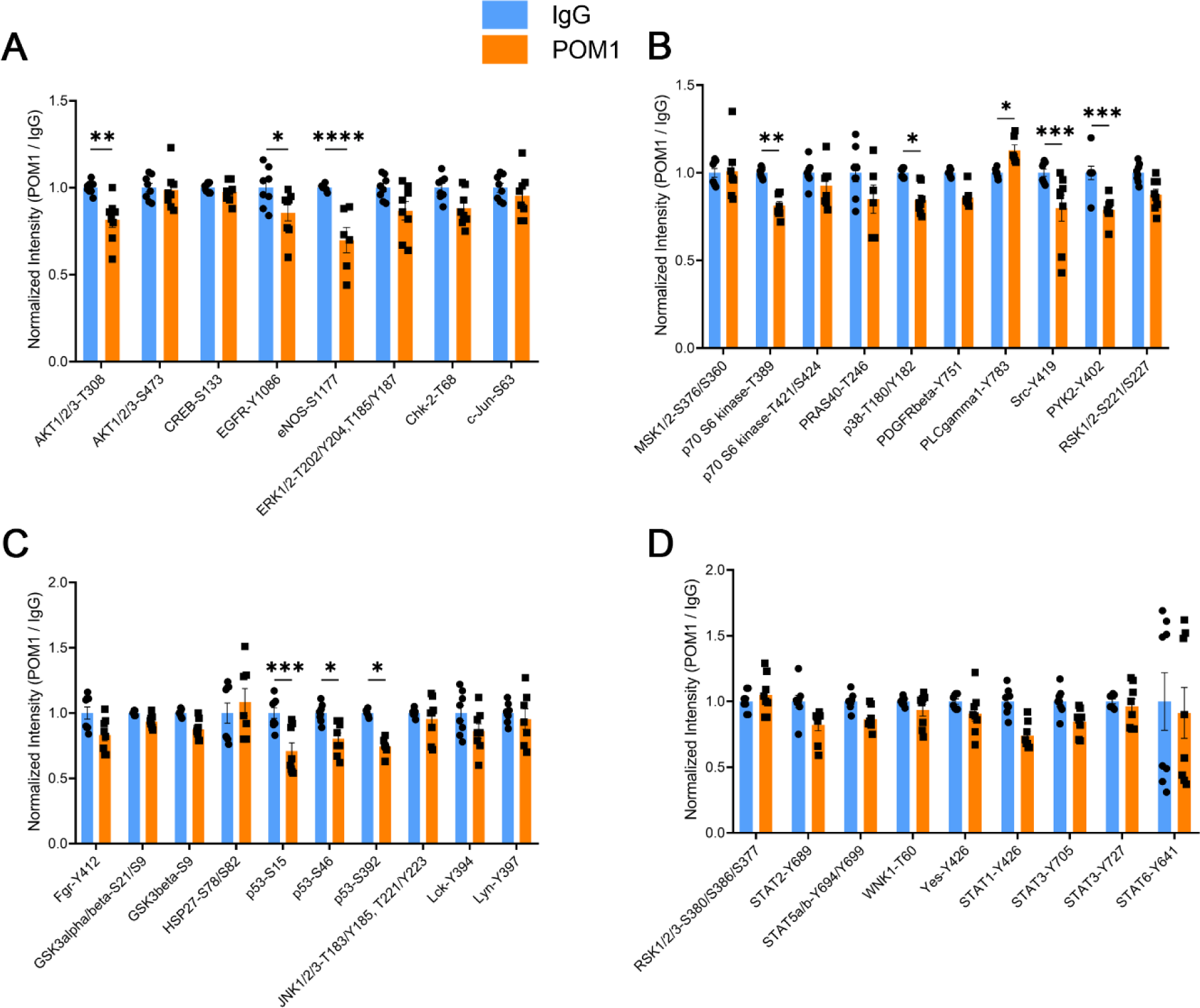
Phospho-kinase array of iNs treated with POM1 or IgG. **(A - D)** Quantification of all 37 phospho-sites in the phospho-kinase array following POM1 or IgG exposure for 2 hours. Data is normalized to the average of the two technical replicates of IgG for each experiment, two-way ANOVA with Sidak’s MCT. *P ≤ 0.05; **P ≤ 0.01; ***P ≤ 0.001; ****P ≤ 0.0001.

## References

1. Terry RD, Masliah E, Salmon DP, Butters N, DeTeresa R, Hill R, et al. Physical basis of cognitive alterations in Alzheimer’s disease: synapse loss is the major correlate of cognitive impairment. Ann Neurol. 1991;30(4):572–80.

2. Targa Dias Anastacio H, Matosin N, and Ooi L. Neuronal hyperexcitability in Alzheimer’s disease: what are the drivers behind this aberrant phenotype? Translational Psychiatry. 2022;12(1):257.

3. Belichenko PV, Brown D, Jeffrey M, and Fraser JR. Dendritic and synaptic alterations of hippocampal pyramidal neurones in scrapie-infected mice. Neuropathology and applied neurobiology. 2000;26(2):143–9.

4. Brown D, Belichenko P, Sales J, Jeffrey M, and Fraser JR. Early loss of dendritic spines in murine scrapie revealed by confocal analysis. Neuroreport. 2001;12(1):179–83.

5. Cunningham C, Deacon R, Wells H, Boche D, Waters S, Diniz CP, et al. Synaptic changes characterize early behavioural signs in the ME7 model of murine prion disease. Eur J Neurosci. 2003;17(10):2147–55.

6. Ferrer I. Synaptic pathology and cell death in the cerebellum in Creutzfeldt-Jakob disease. Cerebellum. 2002;1(3):213–22.

7. Gray BC, Siskova Z, Perry VH, and O’Connor V. Selective presynaptic degeneration in the synaptopathy associated with ME7-induced hippocampal pathology. Neurobiol Dis. 2009;35(1):63–74.

8. Jeffrey M, Halliday WG, Bell J, Johnston AR, MacLeod NK, Ingham C, et al. Synapse loss associated with abnormal PrP precedes neuronal degeneration in the scrapie-infected murine hippocampus. Neuropathol Appl Neurobiol. 2000;26(1):41–54.

9. Siskova Z, Reynolds RA, O’Connor V, and Perry VH. Brain region specific pre-synaptic and post-synaptic degeneration are early components of neuropathology in prion disease. PLoS One. 2013;8(1):e55004.

10. Ojeda-Juárez D, Lawrence JA, Soldau K, Pizzo DP, Wheeler E, Aguilar-Calvo P, et al. Prions induce an early Arc response and a subsequent reduction in mGluR5 in the hippocampus. Neurobiol Dis. 2022;172:105834.

11. Lawrence JA, Aguilar-Calvo P, Ojeda-Juarez D, Khuu H, Soldau K, Pizzo DP, et al. Diminished Neuronal ESCRT-0 Function Exacerbates AMPA Receptor Derangement and Accelerates Prion-Induced Neurodegeneration. J Neurosci. 2023;43(21):3970–84.

12. Sikorska B, Liberski PP, Giraud P, Kopp N, and Brown P. Autophagy is a part of ultrastructural synaptic pathology in Creutzfeldt-Jakob disease: a brain biopsy study. Int J Biochem Cell Biol. 2004;36(12):2563–73.

13. Tao-Cheng JH. Stimulation induces gradual increases in the thickness and curvature of postsynaptic density of hippocampal CA1 neurons in slice cultures. Mol Brain. 2019;12(1):44.

14. Waltereit R, Dammermann B, Wulff P, Scafidi J, Staubli U, Kauselmann G, et al. Arg3.1/Arc mRNA induction by Ca2+ and cAMP requires protein kinase A and mitogen-activated protein kinase/extracellular regulated kinase activation. J Neurosci. 2001;21(15):5484–93.

15. Lyford GL, Yamagata K, Kaufmann WE, Barnes CA, Sanders LK, Copeland NG, et al. Arc, a growth factor and activity-regulated gene, encodes a novel cytoskeleton-associated protein that is enriched in neuronal dendrites. Neuron. 1995;14(2):433–45.

16. Shepherd JD, Rumbaugh G, Wu J, Chowdhury S, Plath N, Kuhl D, et al. Arc/Arg3. 1 mediates homeostatic synaptic scaling of AMPA receptors. Neuron. 2006;52(3):475-84.

17. Chowdhury S, Shepherd JD, Okuno H, Lyford G, Petralia RS, Plath N, et al. Arc/Arg3.1 interacts with the endocytic machinery to regulate AMPA receptor trafficking. Neuron. 2006;52(3):445–59.

18. Zhang W, Wu J, Ward MD, Yang S, Chuang Y-A, Xiao M, et al. Structural basis of arc binding to synaptic proteins: implications for cognitive disease. Neuron. 2015;86(2):490–500.

19. Chen X, Jia B, Araki Y, Liu B, Ye F, Huganir R, et al. Arc weakens synapses by dispersing AMPA receptors from postsynaptic density via modulating PSD phase separation. Cell Res. 2022;32(10):914–30.

20. Okuno H, Minatohara K, and Bito H. Inverse synaptic tagging: An inactive synapse-specific mechanism to capture activity-induced Arc/arg3.1 and to locally regulate spatial distribution of synaptic weights. Semin Cell Dev Biol. 2018;77:43–50.

21. Kraus A, Hoyt F, Schwartz CL, Hansen B, Artikis E, Hughson AG, et al. High-resolution structure and strain comparison of infectious mammalian prions. Mol Cell. 2021;81(21):4540–51.e6.

22. Li Q, Jaroniec CP, and Surewicz WK. Cryo-EM structure of disease-related prion fibrils provides insights into seeding barriers. Nat Struct Mol Biol. 2022;29(10):962–5.

23. Um JW, Kaufman AC, Kostylev M, Heiss JK, Stagi M, Takahashi H, et al. Metabotropic glutamate receptor 5 is a coreceptor for Alzheimer aβ oligomer bound to cellular prion protein. Neuron. 2013;79(5):887–902.

24. Haas LT, Salazar SV, Kostylev MA, Um JW, Kaufman AC, and Strittmatter SM. Metabotropic glutamate receptor 5 couples cellular prion protein to intracellular signalling in Alzheimer’s disease. Brain. 2016;139(Pt 2):526–46.

25. Rösener NS, Gremer L, Wördehoff MM, Kupreichyk T, Etzkorn M, Neudecker P, et al. Clustering of human prion protein and α-synuclein oligomers requires the prion protein N-terminus. Commun Biol. 2020;3(1):365.

26. Um JW, Nygaard HB, Heiss JK, Kostylev MA, Stagi M, Vortmeyer A, et al. Alzheimer amyloid-beta oligomer bound to postsynaptic prion protein activates Fyn to impair neurons. Nat Neurosci. 2012;15(9):1227–35.

27. Fang C, Wu B, Le NTT, Imberdis T, Mercer RCC, and Harris DA. Prions activate a p38 MAPK synaptotoxic signaling pathway. PLoS Pathog. 2018;14(9):e1007283.

28. Schmitt-Ulms G, Legname G, Baldwin MA, Ball HL, Bradon N, Bosque PJ, et al. Binding of neural cell adhesion molecules (N-CAMs) to the cellular prion protein. J Mol Biol. 2001;314(5):1209–25.

29. Khosravani H, Zhang Y, Tsutsui S, Hameed S, Altier C, Hamid J, et al. Prion protein attenuates excitotoxicity by inhibiting NMDA receptors. J Cell Biol. 2008;181(3):551–65.

30. Llorens F, Carulla P, Villa A, Torres JM, Fortes P, Ferrer I, et al. PrPC regulates epidermal growth factor receptor function and cell shape dynamics in Neuro2a cells. J Neurochem. 2013;127(1):124–38.

31. Mantuano E, Azmoon P, Banki MA, Lam MS, Sigurdson CJ, and Gonias SL. A soluble derivative of PrP(C) activates cell-signaling and regulates cell physiology through LRP1 and the NMDA receptor. J Biol Chem. 2020;295(41):14178–88.

32. Foliaki ST, Groveman BR, Yuan J, Walters R, Zhang S, Tesar P, et al. Pathogenic prion protein isoforms are not present in cerebral organoids generated from asymptomatic donors carrying the E200K mutation associated with familial prion disease. Pathogens (Basel, Switzerland). 2020;9(6):482.

33. Foliaki ST, Schwarz B, Groveman BR, Walters RO, Ferreira NC, Orrù CD, et al. Neuronal excitatory-to-inhibitory balance is altered in cerebral organoid models of genetic neurological diseases. Mol Brain. 2021;14:1–23.

34. Foliaki ST, Smith A, Schwarz B, Bohrnsen E, Bosio CM, Williams K, et al. Altered energy metabolism in Fatal Familial Insomnia cerebral organoids is associated with astrogliosis and neuronal dysfunction. PLoS Genet. 2023;19(1):e1010565.

35. Smith A, Groveman BR, Winkler C, Williams K, Walters R, Yuan J, et al. Stress and viral insults do not trigger E200K PrP conversion in human cerebral organoids. PLoS One. 2022;17(10):e0277051.

36. Gojanovich AD, Le NTT, Mercer RCC, Park S, Wu B, Anane A, et al. Abnormal synaptic architecture in iPSC-derived neurons from a multi-generational family with genetic Creutzfeldt-Jakob disease. Stem cell reports. 2024;19(10):1474–88.

37. Matamoros-Angles A, Gayosso LM, Richaud-Patin Y, Di Domenico A, Vergara C, Hervera A, et al. iPS Cell cultures from a Gerstmann-Sträussler-Scheinker patient with the Y218N PRNP mutation recapitulate tau pathology. Mol Neurobiol. 2018;55:3033–48.

38. Groveman BR, Ferreira NC, Foliaki ST, Walters RO, Winkler CW, Race B, et al. Human cerebral organoids as a therapeutic drug screening model for Creutzfeldt–Jakob disease. Sci Rep. 2021;11(1):5165.

39. Groveman BR, Foliaki ST, Orru CD, Zanusso G, Carroll JA, Race B, et al. Sporadic Creutzfeldt-Jakob disease prion infection of human cerebral organoids. Acta neuropathologica communications. 2019;7:1–12.

40. Krejciova Z, Alibhai J, Zhao C, Krencik R, Rzechorzek NM, Ullian EM, et al. Human stem cell-derived astrocytes replicate human prions in a PRNP genotype-dependent manner. J Exp Med. 2017;214(12):3481–95.

41. Herrmann US, Sonati T, Falsig J, Reimann RR, Dametto P, O’Connor T, et al. Prion infections and anti-PrP antibodies trigger converging neurotoxic pathways. PLoS Pathog. 2015;11(2):e1004662.

42. Lakkaraju AKK, Frontzek K, Lemes E, Herrmann U, Losa M, Marpakwar R, et al. Loss of PIKfyve drives the spongiform degeneration in prion diseases. EMBO Mol Med. 2021;13(9):e14714.

43. Tschampa HJ, Kallenberg K, Kretzschmar HA, Meissner B, Knauth M, Urbach H, et al. Pattern of cortical changes in sporadic Creutzfeldt-Jakob disease. AJNR Am J Neuroradiol. 2007;28(6):1114–8.

44. Vitali P, Maccagnano E, Caverzasi E, Henry RG, Haman A, Torres-Chae C, et al. Diffusion-weighted MRI hyperintensity patterns differentiate CJD from other rapid dementias. Neurology. 2011;76(20):1711–9.

45. Baral PK, Wieland B, Swayampakula M, Polymenidou M, Rahman MH, Kav NN, et al. Structural studies on the folded domain of the human prion protein bound to the Fab fragment of the antibody POM1. Acta Crystallogr D Biol Crystallogr. 2012;68(Pt 11):1501–12.

46. Wu B, McDonald AJ, Markham K, Rich CB, McHugh KP, Tatzelt J, et al. The N-terminus of the prion protein is a toxic effector regulated by the C-terminus. eLife. 2017;6.

47. Nuvolone M, Hermann M, Sorce S, Russo G, Tiberi C, Schwarz P, et al. Strictly co-isogenic C57BL/6J-Prnp-/-mice: A rigorous resource for prion science. J Exp Med. 2016;213(3):313–27.

48. Hunter CJ, Remenyi J, Correa SA, Privitera L, Reyskens KMSE, Martin KJ, et al. MSK1 regulates transcriptional induction of Arc/Arg3.1 in response to neurotrophins. FEBS Open Bio. 2017;7(6):821–34.

49. LaCasse RA, Striebel JF, Favara C, Kercher L, and Chesebro B. Role of Erk1/2 activation in prion disease pathogenesis: absence of CCR1 leads to increased Erk1/2 activation and accelerated disease progression. J Neuroimmunol. 2008;196(1-2):16–26.

50. Ferrell JE, and Bhatt RR. Mechanistic Studies of the Dual Phosphorylation of Mitogen-activated Protein Kinase*. J Biol Chem. 1997;272(30):19008–16.

51. Wang C, Ward ME, Chen R, Liu K, Tracy TE, Chen X, et al. Scalable Production of iPSC-Derived Human Neurons to Identify Tau-Lowering Compounds by High-Content Screening. Stem cell reports. 2017;9(4):1221–33.

52. Peng F, Yao H, Bai X, Zhu X, Reiner BC, Beazely M, et al. Platelet-derived Growth Factor-mediated Induction of the Synaptic Plasticity Gene Arc/Arg3.1 *. J Biol Chem. 2010;285(28):21615–24.

53. Lee YJ, Ho SR, Graves JD, Xiao Y, Huang S, and Lin WC. CGRRF1, a growth suppressor, regulates EGFR ubiquitination in breast cancer. Breast Cancer Res. 2019;21(1):134.

54. Ellenrieder V, Zhang JS, Kaczynski J, and Urrutia R. Signaling disrupts mSin3A binding to the Mad1-like Sin3-interacting domain of TIEG2, an Sp1-like repressor. The EMBO Journal. 2002;21(10):2451–60.

55. Buttar NS, DeMars CJ, Lomberk G, Ilyas SI, Bonilla-Velez J, Achra S, et al. Distinct Role of Kruppel-like Factor 11 in the Regulation of Prostaglandin E2 Biosynthesis*. J Biol Chem. 2010;285(15):11433–44.

56. Sorce S, Nuvolone M, Russo G, Chincisan A, Heinzer D, Avar M, et al. Genome-wide transcriptomics identifies an early preclinical signature of prion infection. PLoS Pathog. 2020;16(6):e1008653.

57. Slota JA, Medina SJ, Frost KL, and Booth SA. Neurons and Astrocytes Elicit Brain Region Specific Transcriptional Responses to Prion Disease in the Murine CA1 and Thalamus. Front Neurosci. 2022;16.

58. Liu Y, Guo J, Matoga M, Korotkova M, Jakobsson PJ, and Aguzzi A. NG2 glia protect against prion neurotoxicity by inhibiting microglia-to-neuron prostaglandin E2 signaling. Nat Neurosci. 2024;27(8):1534–44.

59. Balla A, Kim YJ, Varnai P, Szentpetery Z, Knight Z, Shokat KM, et al. Maintenance of Hormone-sensitive Phosphoinositide Pools in the Plasma Membrane Requires Phosphatidylinositol 4-Kinase IIIα. Mol Biol Cell. 2008;19(2):711–21.

60. Shin H-W, Hayashi M, Christoforidis S, Lacas-Gervais S, Hoepfner S, Wenk MR, et al. An enzymatic cascade of Rab5 effectors regulates phosphoinositide turnover in the endocytic pathway. J Cell Biol. 2005;170(4):607–18.

61. Zhang Q, and Fujita M. Why nature evolved GPI-anchored proteins: unique structure characteristics enable versatile cell surface functions. Glycobiology. 2024;34(12).

62. Newton AC, Bootman MD, and Scott JD. Second messengers. Cold Spring Harb Perspect Biol. 2016;8(8):a005926.

63. Korb E, Wilkinson CL, Delgado RN, Lovero KL, and Finkbeiner S. Arc in the nucleus regulates PML-dependent GluA1 transcription and homeostatic plasticity. Nat Neurosci. 2013;16(7):874–83.

64. Tao P, Han X, Wang Q, Wang S, Zhang J, Liu L, et al. A gain-of-function variation in PLCG1 causes a new immune dysregulation disease. Journal of Allergy and Clinical Immunology. 2023;152(5):1292–302.

65. Ferreiro E, Oliveira CR, and Pereira CMF. The release of calcium from the endoplasmic reticulum induced by amyloid-beta and prion peptides activates the mitochondrial apoptotic pathway. Neurobiol Dis. 2008;30(3):331–42.

66. Ruiz A, Matute C, and Alberdi E. Endoplasmic reticulum Ca2+ release through ryanodine and IP3 receptors contributes to neuronal excitotoxicity. Cell Calcium. 2009;46(4):273–81.

67. Rojas M, Yao S, and Lin Y-Z. Controlling Epidermal Growth Factor (EGF)-stimulated Ras Activation in Intact Cells by a Cell-permeable Peptide Mimicking Phosphorylated EGF Receptor *. J Biol Chem. 1996;271(44):27456–61.

68. Groveman BR, Schwarz B, Bohrnsen E, Foliaki ST, Carroll JA, Wood AR, et al. A PrP EGFR signaling axis controls neural stem cell senescence through modulating cellular energy pathways. J Biol Chem. 2023;299(11):105319.

69. Cunningham C, Deacon RM, Chan K, Boche D, Rawlins JN, and Perry VH. Neuropathologically distinct prion strains give rise to similar temporal profiles of behavioral deficits. Neurobiol Dis. 2005;18(2):258–69.

70. Chandler R. Encephalopathy in mice produced by inoculation with scrapie brain material. 1961.

71. Mallucci G, Dickinson A, Linehan J, Klöhn P-C, Brandner S, and Collinge J. Depleting neuronal PrP in prion infection prevents disease and reverses spongiosis. Science (New York, NY). 2003;302(5646):871-4.

72. Mallucci GR, White MD, Farmer M, Dickinson A, Khatun H, Powell AD, et al. Targeting cellular prion protein reverses early cognitive deficits and neurophysiological dysfunction in prion-infected mice. Neuron. 2007;53(3):325–35.

73. Kawata M, McClatchy DB, Diedrich JK, Olmer M, Johnson KA, Yates JR, et al. Mocetinostat activates Krüppel-like factor 4 and protects against tissue destruction and inflammation in osteoarthritis. JCI insight. 2023;8(17).

74. Zecha J, Satpathy S, Kanashova T, Avanessian SC, Kane MH, Clauser KR, et al. TMT Labeling for the Masses: A Robust and Cost-efficient, In-solution Labeling Approach. Mol Cell Proteomics. 2019;18(7):1468–78.

75. Polymenidou M, Moos R, Scott M, Sigurdson C, Shi YZ, Yajima B, et al. The POM monoclonals: a comprehensive set of antibodies to non-overlapping prion protein epitopes. PLoS One. 2008;3(12):e3872.

